# viGEN: An open source pipeline for the detection and quantification of viral RNA in human tumors

**DOI:** 10.1101/099788

**Authors:** Krithika Bhuvaneshwar, Lei Song, Subha Madhavan, Yuriy Gusev

## Abstract

An estimated 17% of cancers worldwide are associated with infectious causes. The extent and biological significance of viral presence/infection in actual tumor samples is generally unknown but could be measured using human transcriptome (RNA-seq) data from tumor samples.

We present an open source bioinformatics pipeline viGEN, which combines existing well-known and novel RNA-seq tools for not only the detection and quantification of viral RNA, but also variants in the viral transcripts.

The pipeline includes 4 major modules: The first module allows to align and filter out human RNA sequences; the second module maps and count (remaining un-aligned) reads against reference genomes of all known and sequenced human viruses; the third module quantifies read counts at the individual viral genes level thus allowing for downstream differential expression analysis of viral genes between experimental and controls groups. The fourth module calls variants in these viruses. To the best of our knowledge, there are no publicly available pipelines or packages that would provide this type of complete analysis in one open source package.

In this paper, we applied the viGEN pipeline to two case studies. We first demonstrate the working of our pipeline on a large public dataset, the TCGA cervical cancer cohort. We also performed additional in-depth analyses on a small focused study of TCGA liver cancer patients. In this cohort, we perform viral-gene quantification, viral-variant extraction and survival analysis. This allowed us to find differentially expressed viral-transcripts and viral-variants between the groups of patients, and connect them to clinical outcome.

From our analyses, we show that we were able to successfully detect the human papilloma virus among the TCGA cervical cancer patients. We compared the viGEN pipeline with two metagenomics tools and demonstrate similar sensitivity/specificity. We were also able to quantify viral-transcripts and extract viral-variants using the liver cancer dataset. The results presented corresponded with published literature in terms of rate of detection, viral gene expression patterns and impact of several known variants of HBV genome. Results also show novel information about distinct patterns of expression and co-expression in Hepatitis B and the Human Endogenous Retrovirus (HERV) K113 viruses.

This pipeline is generalizable, and can be used to provide novel biological insights into the significance of viral and other microbial infections in complex diseases, tumorigeneses and cancer immunology. The source code, with example data and tutorial is available at: https://github.com/ICBI/viGEN/.

## INTRODUCTION

An estimated 17% of cancers worldwide are associated with infectious causes. These infectious agents include viruses, bacteria, parasites and other microbes. Examples of viruses include human papilloma viruses (HPVs) in cervical cancer, epstein-Barr virus (EBV) in nasopharyngeal cancer, hepatitis B and C in liver cancer (HBV and HCV), human herpes virus 8 (HHV-8) in Kaposi sarcoma (KS); human T-lymphotrophic virus-1 (HTLV-1) in adult T cell lymphocytic leukemia (ATL) and non-Hodgkin lymphoma; merkel cell polyomavirus (MCV) in Merkel cell carcinoma [1]. Bacteria such as Helicobacter pylori have been implicated in stomach cancer. Parasites have also been associated with cancer, examples are Opisthorchis viverrini and Clonorchis sinensis in bile duct cancer and Schistosoma haematobium in bladder cancer [1]. Detection and characterization of these infectious agents in tumor samples can give us better insights into disease mechanisms and their treatment [2].

Vaccines have been developed to help protect against infection from the many cancers. But these vaccines can only be used to help prevent infection and cannot treat existing infections [1]. There are several screening methods widely used to detect viral infections, especially for blood borne viruses including HBV, HCV, HIV and HTLV. These include the enzyme linked immunosorbent assay (ELISA or EIA) [3], chemluminescent immunoassay (ChLIA), Indirect fluorescent antibody (IFA), Western blot (WB), Polymerase Chain Reaction (PCR), and Rapid immunoassays [4]. ELISA and WB test detects and measures antibodies in serum taken from the patient’s blood, and are typically prescribed after certain symptoms are observed in the patient.

There are several challenges in detection of viruses in tumors including loss of viral information in progressed tumors and limited or latent replication resulting in low transcription of tumors [5]. The extent and biological significance of viral presence/infection in actual tumor samples is generally unknown but could be measured using human transcriptome data from tumor samples.

The popularity of next-generation sequencing (NGS) technology has exploded in the last decade. NGS technologies are able to perform rapid sequencing, and in a massively parallel fashion [6]. In recent years, applications of NGS technologies in clinical diagnostics have been on the rise [7, 8]. Amongst the various NGS technologies, whole-transcriptome sequencing, also called RNA-seq has been very popular with methods and tools being actively developed. Exploring the genome using RNA-seq gives a different insight than looking at the DNA since the RNA-seq would have captured actively transcribed regions. Every aspect of data output from this technology is now being used for research, including detection of viruses and bacteria [9-11]. They are also independent of prior sequence information, and require less starting material compared to conventional cloning based methods, making it a powerful and exciting new technology in virology [6]. These high throughput technologies give us direct evidence of infection in the tissue as compared to ELISA-based assays, which only proves presence of infection somewhere in the human body. RNA-seq technology has hence enabled the exploration and detection of viral infections in human tumor samples. This technology also enables detection of variants in viral genome, which have been connected to clinical outcome [12] [13].

In recent years, US regulators approved a viral based cancer therapy [14], proving that the study of viruses in the human transcriptome has biomedical interest, and is paving the way for promising research and new opportunities.

In this paper, we present our pipeline viGEN to not only detect and quantify read counts at the individual viral genes level, but also detect viral variants from human RNA-seq data. The characterization of viral variants helps enable better epidemiological analysis. The input file to our pipeline is a fastq [15] file, so our viGEN pipeline can be extended to work with genomic data from any NGS technology. Our pipeline can also be used to detect and explore not only viruses, but other microbes as well, as long as the sequence information is available in NCBI [16].

We applied our viGEN pipeline to two case studies as a proof of concept - a dataset of 304 cervical cancer patients, and a set of 50 liver cancer patients, both from the TCGA collection. We first applied the pipeline to the transcriptome of cervical cancer patients to see if we are able to detect the human papilloma viruses. We also performed additional in-depth analyses on a small focused study of liver cancer patients. In this cohort, we perform viral-gene quantification, viral-variant extraction and survival analysis.

From our analyses, we show that we were able to successfully detect the human papilloma virus among the TCGA cervical cancer patients. We compared the viGEN pipeline with two metagenomics tools and demonstrate similar sensitivity/specificity. We were also able to quantify viral-transcripts and extract viral-variants using the liver cancer dataset. This enabled us to perform downstream analysis to give us new insights into disease mechanisms.

In addition to the two case studies, we have made available an end-to-end tutorial demonstrated on a publicly available RNA-seq sample from an HBV liver cancer patient from NCBI SRA (http://www.ncbi.nlm.nih.gov/bioproject/PRJNA279878). We also provided step-by-step instructions on how to run our viGEN pipeline on this sample data, along with the code at https://github.com/ICBI/viGEN/ and demonstrate the detection of HBV transcripts in this sample. This allows other users to apply this pipeline to explore viruses in their data and disease of interest. We are currently implementing the viGEN pipeline in the Seven Bridges Cancer Genomics Cloud [17].

There are a number of existing pipelines that detect viruses from human transcriptome data. Of these, very few pipelines offer quantification at the gene expression level. A comprehensive comparison of these pipelines is provided in Table 1. Our goal was not to compete with these other tools, but to offer a convenient and complete end–to-end publicly available pipeline to the bioinformatics community. To the best of our knowledge there are no publicly available pipelines or packages that would provide this type of complete analysis in one package. Customized solutions have been reported in the literature however were not made public.

**Table 1.** Comparison of existing pipelines that detect viruses from human transcriptome data.

| **Tool Name** | **Detect viruses from Human RNA-seq data** | **Perform quantification at viral-gene/CDS level** | **Works on DNaseq, RNAseq or Both** | **Variant calling at viral-variant level** | **Discover viral integration sites** | **Other comments** |
| --- | --- | --- | --- | --- | --- | --- |
| Virana [1] | Yes | Identifies microbial transcripts, does not quantify | Both | No | Yes | Also offers analysis of homologs |
| VirusSeq [2] | Yes | No | Both | No | Yes | Pre-designed to work on a select set of 18 viruses |
| Viral Fusion Seq [3] | Yes | No | Both | No | Yes | Can also detect fusion events |
| Virus Finder [4] | Yes | No | Both | No | Yes | Can be applied to samples infected with undiagnosed viruses |
| PathSeq [5] | Yes | No | Both | No | No |  |
| RINS [6] | Yes | No | RNA-seq | No | No | Generates contigs with these non-human sequences |
| Kraken [7] | Yes | No | No | No | No | Metagenomic analysis tool |
| Cetrifuge [8] | Yes | No | No | No | No | Metagenomic analysis tool |
| viGEN (our pipeline) | Yes | Yes | Both | Yes | No |  |

In the future, our plan is to package this pipeline and make it available to users through Bioconductor [18], allowing users to perform analysis on either their local computer or the cloud.

## MATERIALS AND METHODS

In this paper, we applied our viGEN pipeline to two case studies as a proof of concept - a dataset of 304 cervical cancer patients, and a set of 50 liver cancer patients, both from the TCGA collection [19]. We first applied the pipeline to the transcriptome of cervical cancer patients to see if we are able to detect the human papilloma viruses. We also performed additional in-depth analyses on a small focused study of liver cancer patients afflicted with Hepatitis B virus. In this cohort, we perform viral-gene quantification, viral-variant extraction and survival analysis. The results from these analyses allowed us to compare experimental and control groups using viral-gene expression data and viral-variant data, and give us insights into their impacts on the tumor, and disease mechanisms.

In the following sections, we describe the viGEN pipeline, and the two case studies.

### The viGEN pipeline

The viGEN pipeline includes 4 major modules. Figure 1 shows an image of our viGEN pipeline.

**Figure 1:**
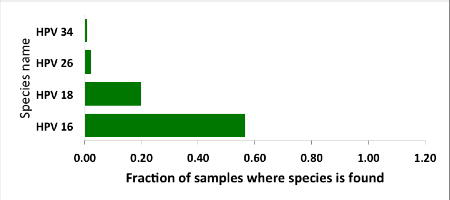
viGEN pipeline. Each module has a color, shown in the legend.

### Module 1: Viral genome level analysis (filtered human sample input)

In Module 1 (labelled as ‘filtered human sample input’), the human RNA sequences were aligned to the human-reference genome using RSEM [20] tool. One of the outputs of RSEM includes sequences that did not align to the human genome (hence the name ‘filtered human sample input’). These un-aligned sequences were taken and aligned to the viral reference file using popular alignment tools BWA [21] and Bowtie2 [22].

### Module 2: Viral genome level analysis (unfiltered human sample input)

In Module 2 (labelled as ‘unfiltered human sample input’), the human RNA seq sequences were directly aligned to the viral reference using Bowtie2 without any filtering. The reason for using two methods to obtain the viral genomes in human RNA-seq data (Module 1 and Module 2) was to allow us to be as comprehensive as possible in viral detection.

The aligned reads from Module 1 and 2 were in the form of BAM files [23], from which read counts were obtained for each viral genome species (referred to as ‘genome level counts’) using Samtools idxstats [24] or Picard BAMIndexStats [25] tools. Using the genome level counts, we estimated the number of reads that covered the genome, a form of viral copy number. Viral copy number was defined as in equation below:

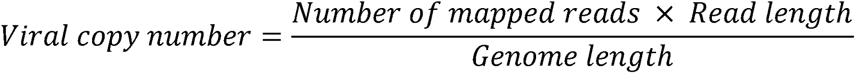

Only those viral species with copy number more than a threshold are selected for the next module.

### Module 3: Viral gene expression analysis

The BAM files from Module 1 and 2 (from Bowtie2 and BWA) were input into the Module 3 (referred to as ‘viral gene expression level analysis’), which calculated quantitate read counts at the individual viral genes level. We found existing RNA-seq quantification tools to be not sensitive enough for viruses, and hence developed our own algorithm for this module. Our in-house algorithm used region-based information from the general-feature-format (GFF) files [26] of each viral genome, and the reads from the BAM file. It created a summary file, which had a total count of reads within or on the boundary of each region in the GFF file. This is repeated for each sample and for each viral GFF file. At the end, a matrix is obtained where the features (rows) are regions from the GFF file, and the columns are samples. The read count output from Module 3 (viral gene expression module) allowed for downstream differential expression analysis of viral genes between experimental and controls groups. The source code for our in-house algorithm written using the R programming language [27] has been made public at available at github.com/ICBI/viGEN.

### Module 4: Viral RNA variant calling module

The BAM files from Module 1 and 2 (from Bowtie2) were also input to Module 4 to detect mutations in the transcripts from these viruses (referred to as ‘viral RNA variant calling module’). The BAM files were first sorted coordinate-wise using Samtools [24]; PCR duplicates were removed using tool Picard [25], then the chromosomes in the BAM file were ordered in the same way as the reference file using Picard. The Viral reference file was created from combining all known and sequenced human viruses obtained from NCBI [16]. Because viral variants are known to be low frequency, we have selected a variant calling tool Varscan2 [28], which allows detection of low-frequency variants [29]. Low quality and low depth variants were flagged, but not filtered out, in case these low values were attributed to low viral load. Once the variants were obtained, they were merged to form a multi-sample VCF file. Only variants that had a variant in at-least one sample were retained. PLINK [30] was used to perform case-control association test (Fishers Exact Test) to compare groups.

## Tutorial in Github

The viGEN pipeline is easy to implement because our pipeline incorporates existing best practices and tools available. For Module 3, we found existing RNA-seq quantification tools to be not sensitive enough for viruses, and hence developed our own algorithm. The source code for the in-house algorithm, along with a tutorial on how to execute the code on sample data has been made public at https://github.com/ICBI/viGEN/.

Since access to TCGA raw data is controlled access, we could not use this dataset to create a publicly available tutorial. So we used a publicly available RNA-seq dataset to demonstrate our pipeline with an end-to-end workflow. We chose one sample (SRR1946637) from publicly available HBV liver cancer RNA-seq dataset from NCBI SRA (http://www.ncbi.nlm.nih.gov/bioproject/PRJNA279878). This dataset is also available through EBI SRA (http://www.ebi.ac.uk/ena/data/view/SRR1946637). The dataset consists of 50 HBV Liver cancer patients, and 5 adjacent normal liver tissues. We downloaded the raw reads for one sample, and applied our viGEN pipeline to it and were able to successfully detect HBV transcripts in this sample. A step-by-step workflow that includes – description of tools, code, intermediate and final analysis results are provided in Github: https://github.com/ICBI/viGEN/. This tutorial has also been provided as Additional File 1.

## Custom reference index

We were interested in exploring all viruses existing in humans. So we first obtained reference genomes of all known and sequenced human viruses obtained from NCBI [16] (745 viruses) and merged them into one file (referred to as the ‘viral reference file’) in fasta file format [31]. This file has been shared in our Github page.

## Case studies

### Cervical cancer dataset

Cervical cancer is caused by the Human Papilloma Virus (HPV). This dataset consisted of 304 cervical cancer patients in the TCGA data collection. These samples were primary tumors from either Cervical Squamous Cell Carcinoma or Endocervical Adenocarcinoma where RNA-seq data was available.

We applied our viGEN pipeline on these samples using the Seven Bridges platform (https://cgc.sbgenomics.com). Among the 304 cervical cancer patients, 22 patients had virus detection confirmed by PCR or other lab methods and made available through the clinical data. So we used this information from the 22 patients to estimate the sensitivity and specificity of our viGEN pipeline.

### Liver cancer dataset

This dataset consisted of 50 liver cancer patients in the TCGA data collection. 25 of these patients were afflicted with Hepatitis B virus (labelled ‘HepB’), while the rest of the 25 patients had a co-infection of both Hepatitis B and C viruses (labelled ‘HepB+C’). Information about viral presence was obtained from ‘Viral Hepatitis Serology’ attribute from the clinical information.

We first applied the viGEN pipeline on the 50 samples, using the Globus Genomics platform [32]. Once the viral genomes were detected, we then chose only the high abundance viral species for the gene quantification step and viral variant detection steps (Module 3 and 4 respectively).

We then performed a focused analysis on this dataset. We used the viral-gene expression read counts, to examine the differences between “Dead” and “Alive” samples. The analysis was performed using a Bioconductor software package called EdgeR [33] in the R programming language (http://www.R-project.org). Cox proportional hazards (Cox PH) regression model [34] was applied to this group to look at the association of viral-gene expression data with overall survival. We also compared the dead and alive samples at the viral RNA variant level using a tool called PLINK to see if it can add valuable information to the tumor landscape in humans.

## RESULTS

### Detection of HPV in cervical cancer patients

We used our viGEN pipeline to detect viruses in the RNA of human cervical tissue and obtained viral copy number for each species. We used a threshold copy number of 10 as a ‘positive’ viral detection for both HPV-16, HPV-18 and HPV-26 viruses. Based on this criterion, HPV-16 was detected in 53% of the samples, HPV-18 in 13% of the samples and HPV-26 in 0.3 % of the samples (Figure 2).

**Figure 2:**
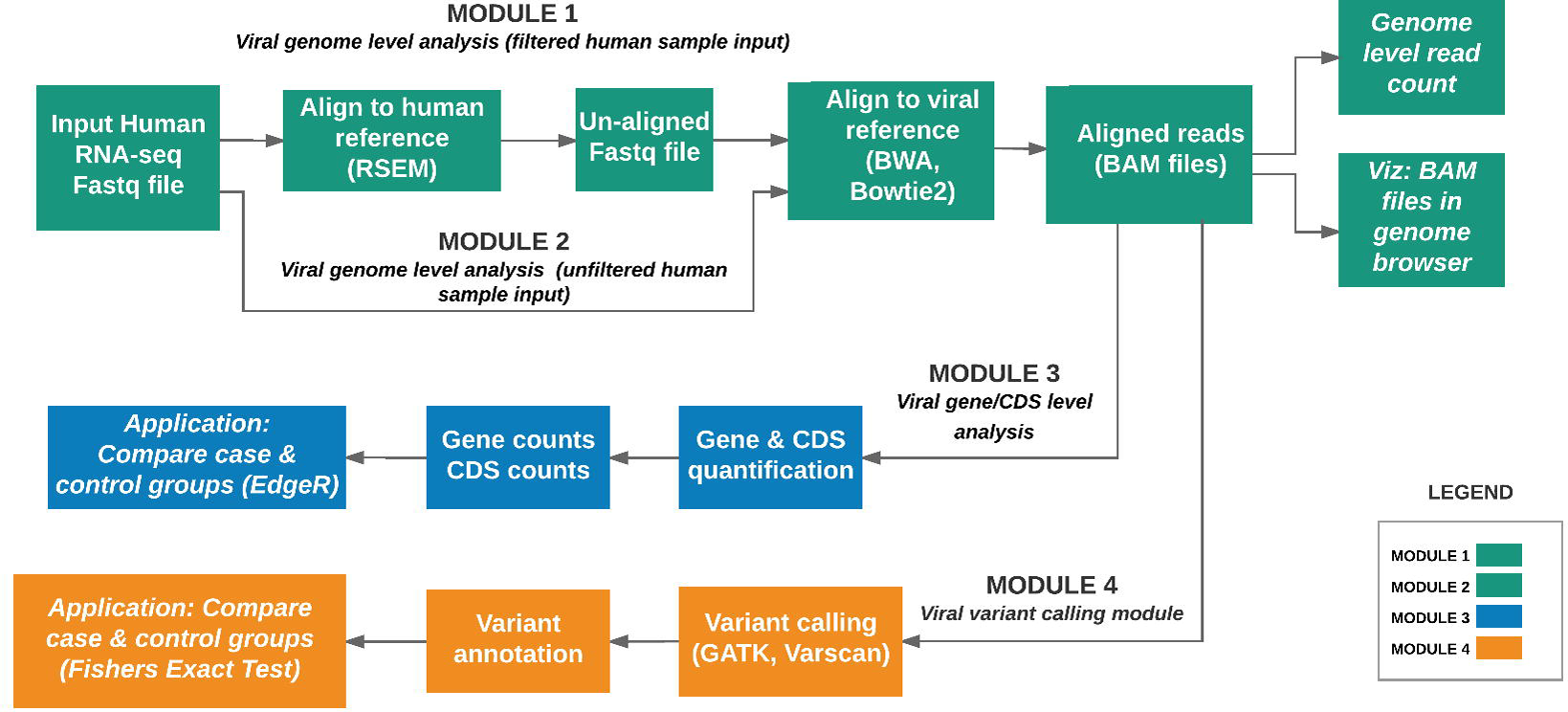
The HPV viruses detected in cervical cancer patients using the viGEN pipeline.

We obtained the clinical data for this TCGA cervical cancer cohort from the cBio portal [35]. Among the 304 patients, 22 patients had virus detection confirmed by PCR or other lab methods and made available through the clinical data. Out of the 22 patients, 12 patients had the HPV-16 virus, 4 patients had HPV-18, and the rest had other HPV viruses. So we used this information from the clinical data to estimate the sensitivity and specificity of our viGEN pipeline. We got a sensitivity of 83% and specificity of 60% for HPV-16 detection (Table 2 A); and a sensitivity of 75% and specificity of 94% for HPV-18 detection (Table 2 B)

**Table 2(A):** Estimation of sensitivity and specificity for HPV-16 detection in TCGA cervical cancer samples using the viGEN pipeline.

| PCR or other lab techniques for<br>screening viruses (obtained from<br>clinical data) |  |  |  |  |
| --- | --- | --- | --- | --- |
|  |  | Positive | Negative | Total |
| RNA-seq from<br>tumor tissue<br>(output of<br>algorithm) | Positive | 10 | 4 | 14 |
|  | Negative | 2 | 6 | 8 |
|  | Total | 12 | 10 | 22 |

**Table 2(B):** Estimation of sensitivity and specificity for HPV-18 detection in TCGA cervical cancer samples using the viGEN pipeline.

| PCR or other lab techniques for<br>screening viruses (obtained from<br>clinical data) |  |  |  |  |
| --- | --- | --- | --- | --- |
|  |  | Positive | Negative | Total |
| RNA-seq from<br>tumor tissue<br>(output of<br>algorithm) | Positive | 3 | 1 | 4 |
|  | Negative | 1 | 17 | 18 |
|  | Total | 4 | 18 | 22 |

## Additional analysis in liver cancer patients

### Detection of Hepatitis B virus at the genome level

We applied our viGEN pipeline (modules 1 and 2) on the RNAseq data from the TCGA liver cancer tumors, and obtained genome-level read counts for each viral species. We used a threshold copy number of 10 to define a positive detection of the Hepatitis B virus.

Once the viral genomes were detected, we short-listed the high abundance viral species for the viral-gene quantification step and viral-variant detection steps (Module 3 and 4 respectively). High abundance was defined as those virus species that were detected in at-least 5 samples. In addition to Hepatitis B and C viruses, several other viruses came up in this short list including Human endogenous retrovirus K113 (HERV K113 and others. A complete list is provided in Table 3.

**Table 3:** Viruses species detected in at-least 5 samples.

| **Name of Virus** |
| --- |
| Hepatitis C virus |
| Tick-borne encephalitis virus |
| Hepatitis C virus genotype 2 |
| Cutthroat trout virus |
| Human endogenous retrovirus K113 |
| Lunk virus NKS-1 |
| Hepatitis C virus genotype 6 |
| Hepatitis B virus |

### Comparing dead and alive samples in the using viral gene expression data

To get a more detailed overview of the viral landscape, we applied Module 3 of the viGEN pipeline to the liver cancer dataset. This allowed us to quantify viral-gene expression regions in the RNA of liver tumor tissues. We then used those results to examine the differences between dead and alive samples.

Out of 25 HepB patients, 16 were alive (baseline group), and 9 dead (comparison group) as per the clinical data. It is known that these patients were afflicted with the Hepatitis B virus and hence many of the differentially expressed regions were from this viral genome. But as we know, other viruses also coexist in humans. This was confirmed by the presence of differentially expressed viral-regions from other viruses.

The differentially expressed regions that were significant among the results are shown in Table 4 (A) and Table 4 (B). Table 4 (A) lists only the differentially expressed regions from Hepatitis B virus and Table 4 (B) shows the differentially expressed regions from other viruses.

**Table 4:**
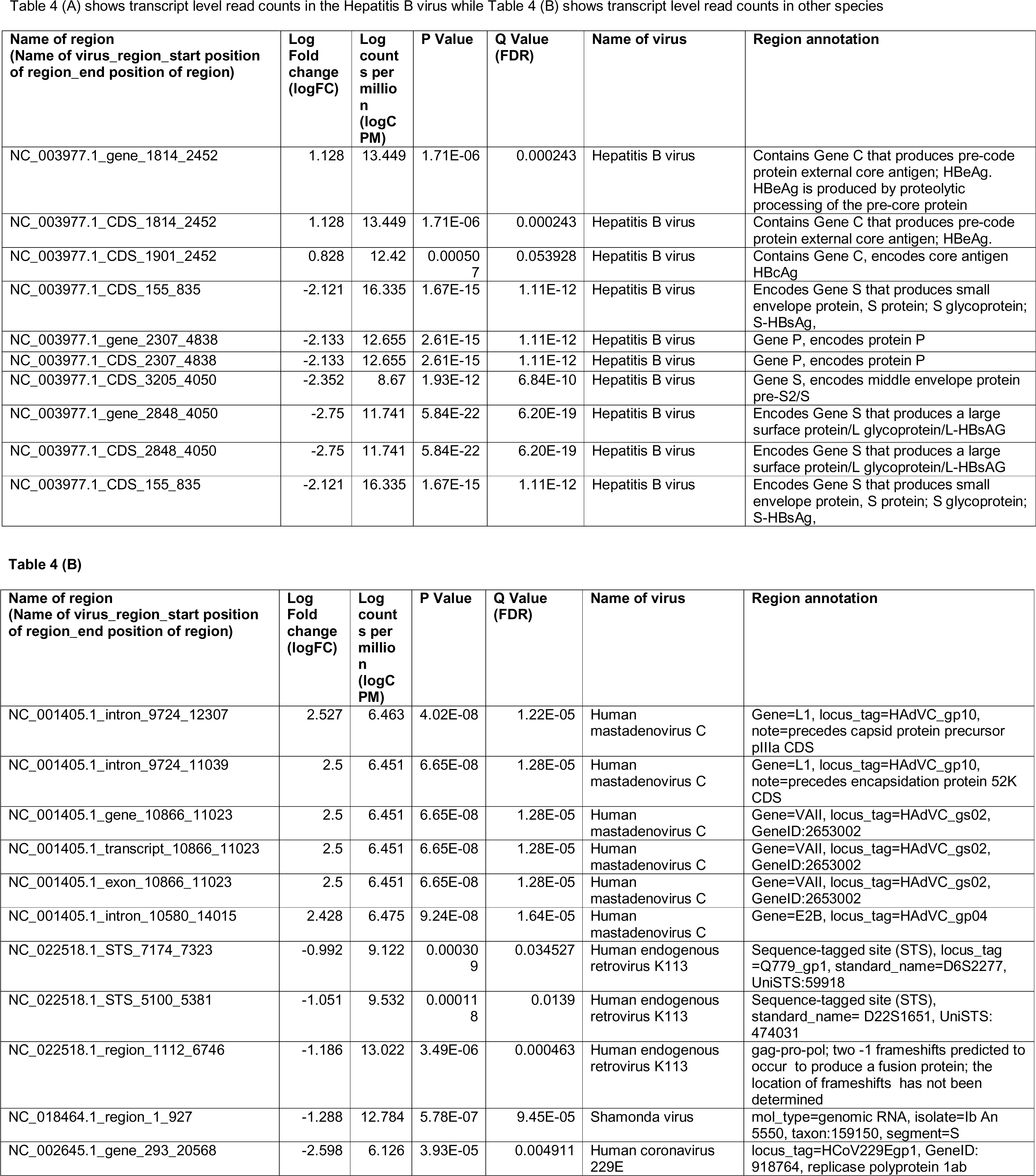
Differential expression analysis of transcript level read counts Liver cancer dataset comparing Dead and Alive samples. These results shown used the viral-gene data obtained from Module 1 (using alignment tool Bowtie2) + Module 3. The table shows results with q value < 0.06 and sorted based on LogFC in the descending order.

| Name of region<br>(Name of virus_region_start position<br>of region_end position of region) | Log<br>Fold<br>change<br>(logFC) | Log<br>count<br>s per<br>millio<br>n<br>(logC<br>PM) | P Value | Q Value<br>(FDR) | Name of virus | Region annotation |
| --- | --- | --- | --- | --- | --- | --- |
| NC_003977.1_gene_1814_2452 | 1.128 | 13.449 | 1.71E-06 | 0.000243 | Hepatitis B virus | Contains Gene C that produces pre-code protein external core antigen; HBeAg. HBeAg is produced by proteolytic processing of the pre-core protein |
| NC_003977.1_CDS_1814_2452 | 1.128 | 13.449 | 1.71E-06 | 0.000243 | Hepatitis B virus | Contains Gene C that produces pre-code protein external core antigen; HBeAg. |
| NC_003977.1_CDS_1901_2452 | 0.828 | 12.42 | 0.000507 | 0.053928 | Hepatitis B virus | Contains Gene C, encodes core antigen HBcAg |
| NC_003977.1_CDS_155_835 | -2.121 | 16.335 | 1.67E-15 | 1.11E-12 | Hepatitis B virus | Encodes Gene S that produces small envelope protein, S protein; S glycoprotein; S-HBsAg, |
| NC_003977.1_gene_2307_4838 | -2.133 | 12.655 | 2.61E-15 | 1.11E-12 | Hepatitis B virus | Gene P, encodes protein P |
| NC_003977.1_CDS_2307_4838 | -2.133 | 12.655 | 2.61E-15 | 1.11E-12 | Hepatitis B virus | Gene P, encodes protein P |
| NC_003977.1_CDS_3205_4050 | -2.352 | 8.67 | 1.93E-12 | 6.84E-10 | Hepatitis B virus | Gene S, encodes middle envelope protein pre-S2/S |
| NC_003977.1_gene_2848_4050 | -2.75 | 11.741 | 5.84E-22 | 6.20E-19 | Hepatitis B virus | Encodes Gene S that produces a large surface protein/L glycoprotein/L-HBsAG |
| NC_003977.1_CDS_2848_4050 | -2.75 | 11.741 | 5.84E-22 | 6.20E-19 | Hepatitis B virus | Encodes Gene S that produces a large surface protein/L glycoprotein/L-HBsAG |
| NC_003977.1_CDS_155_835 | -2.121 | 16.335 | 1.67E-15 | 1.11E-12 | Hepatitis B virus | Encodes Gene S that produces small envelope protein, S protein; S glycoprotein; S-HBsAg, |

| Name of region<br>(Name of virus_region_start position<br>of region_end position of region) | Log<br>Fold<br>change<br>(logFC) | Log<br>count<br>s per<br>millio<br>n<br>(logC<br>PM) | P Value | Q Value<br>(FDR) | Name of virus | Region annotation |
| --- | --- | --- | --- | --- | --- | --- |
| NC_001405.1_intron_9724_12307 | 2.527 | 6.463 | 4.02E-08 | 1.22E-05 | Human<br>mastadenovirus C | Gene=L1, locus_tag=HAdVC_gp10,<br>note=precedes capsid protein precursor<br>pIIIa CDS |
| NC_001405.1_intron_9724_11039 | 2.5 | 6.451 | 6.65E-08 | 1.28E-05 | Human<br>mastadenovirus C | Gene=L1, locus_tag=HAdVC_gp10,<br>note=precedes encapsidation protein 52K<br>CDS |
| NC_001405.1_gene_10866_11023 | 2.5 | 6.451 | 6.65E-08 | 1.28E-05 | Human<br>mastadenovirus C | Gene=VAII, locus_tag=HAdVC_gs02,<br>GeneID:2653002 |
| NC_001405.1_transcript_10866_11023 | 2.5 | 6.451 | 6.65E-08 | 1.28E-05 | Human<br>mastadenovirus C | Gene=VAII, locus_tag=HAdVC_gs02,<br>GeneID:2653002 |
| NC_001405.1_exon_10866_11023 | 2.5 | 6.451 | 6.65E-08 | 1.28E-05 | Human<br>mastadenovirus C | Gene=VAII, locus_tag=HAdVC_gs02,<br>GeneID:2653002 |
| NC_001405.1_intron_10580_14015 | 2.428 | 6.475 | 9.24E-08 | 1.64E-05 | Human<br>mastadenovirus C | Gene=E2B, locus_tag=HAdVC_gp04 |
| NC_022518.1_STS_7174_7323 | -0.992 | 9.122 | 0.00030<br>9 | 0.034527 | Human endogenous<br>retrovirus K113 | Sequence-tagged site (STS), locus_tag<br>=Q779_gp1, standard_name=D6S2277,<br>UniSTS:59918 |
| NC_022518.1_STS_5100_5381 | -1.051 | 9.532 | 0.00011<br>8 | 0.0139 | Human endogenous<br>retrovirus K113 | Sequence-tagged site (STS),<br>standard_name= D22S1651, UniSTS:<br>474031 |
| NC_022518.1_region_1112_6746 | -1.186 | 13.022 | 3.49E-06 | 0.000463 | Human endogenous<br>retrovirus K113 | gag-pro-pol; two -1 frameshifts predicted to<br>occur to produce a fusion protein; the<br>location of frameshifts has not been<br>determined |
| NC_018464.1_region_1_927 | -1.288 | 12.784 | 5.78E-07 | 9.45E-05 | Shamonda virus | mol_type=genomic RNA, isolate=Ib An<br>5550, taxon:159150, segment=S |
| NC_002645.1_gene_293_20568 | -2.598 | 6.126 | 3.93E-05 | 0.004911 | Human coronavirus<br>229E | locus_tag=HCoV229Egp1, GeneID:<br>918764, replicase polyprotein 1ab |

From the differential expression analyses, the two most informative results were (1) a region of the Hepatitis B genome that produced the HBeAg and HBcAg proteins were overexpressed in the dead patients and (2) another region of the Hepatitis B genome that produced HBsAg protein was overexpressed in the alive patients.

In detail, we saw several important findings as described below:

a. Region NC_003977.1_CDS_1814_2452 of the Hepatitis B genome was 2.18 times overexpressed (log fold change = +1.128) in dead patients. This region contains Gene C that produces pre-code protein external core antigen; HBeAg. HBeAg is produced by proteolytic processing of the pre-core protein
b. Region NC_003977.1_CDS_1901_2452 which was 1.74 times overexpressed (log fold change = +0.8, FDR = 0.053) in dead patients contains Gene C as above, but encodes a different core antigen HBcAg
c. Region NC_003977.1_CDS_2848_4050 of the Hepatitis B genome was 6.73 times over expressed (log fold change = -2.7) in the alive patients of compared to the dead’patients. This region encodes Gene S that produces a large surface protein/L glycoprotein/L-HBsAG
d. We also found several regions of the Human endogenous retrovirus K113 (HERV K113) viral genome (NC_022518.1_region_1112_6746, NC_022518.1_STS_5100_5381 and NC_022518.1_STS_7174_7323) to be about 2 times overexpressed on average in alive patients (log fold change = -1.186, -1.051, -0.992).

### Survival analysis (Cox Regression) using viral gene expression data

Based on the results from previous section, we selected two most informative regions from the Hepatitis B genome (log counts per million from NC_003977.1_CDS_2848_4050, NC_003977.1_CDS_1814_2452) for a Cox Proportional Hazard (Cox PH) model to look at association with overall survival event and time. This model was applied on the 25 Hep B and 25 HepB+HepC samples to maximize power. The result from this model (Table 5), are consistent with the results from differential expression analysis:

a. The Cox PH model shows that assuming other covariant to be constant, unit increase in expression of this region NC_003977.1_CDS_1814_2452, increases the hazard of event (death) by 70%.
b. On the other hand, that assuming other covariant to be constant, unit increase in expression of this region NC_003977.1_CDS_2848_4050, decreases the hazard of event (death) by 43%.
c. The overall model is significant with p-value < 0.05 from the Log rank test (also called Score test).

**Table 5:** Cox proportional hazard survival analysis (across 25 HepB samples and 25 HepB + HepC Samples). These results shown used the viral-gene expression data obtained from Module 1 (using alignment tool Bowtie2) + Module 3. *Coef: coefficient (Beta) of the model; exp(coef): Hazard Ratio; se(coef): Standard Error; Pr(<|z|): P-value*

| **Formula:** |  |  |  |  |  |
| --- | --- | --- | --- | --- | --- |
| coxph(formula = survObject ~ NC_003977.1_CDS_2848_4050 + NC_003977.1_CDS_1814_2452) |  |  |  |  |  |
| **Results from the model:** |  |  |  |  |  |
| n= 37, number of events= 5 |  |  |  |  |  |
| (13 observations deleted due to missingness) |  |  |  |  |  |
| Covariate | coef | exp(coef) | se(coef) | Z | Pr(>\|z\|) |
| NC_003977.1_CDS_2848_4050 | -0.5548 | 0.5742 | 0.7434 | -0.746 | 0.456 |
| NC_003977.1_CDS_1814_2452 | 0.5302 | 1.6993 | 0.6145 | 0.863 | 0.388 |
| Covariate | exp(coef ) | exp(-coef) | Lower 0.95 | Upper 0.95 |  |
| NC_003977.1_CDS_2848_4050 | 0.5742 | 1.7415 | 0.1337 | 2.465 |  |
| NC_003977.1_CDS_1814_2452 | 1.6993 | 0.5885 | 0.5096 | 5.667 |  |
| Concordance= 0.654 (se = 0.188 ) |  |  |  |  |  |
| Rsquare= 0.12 (max possible= 0.329 ) |  |  |  |  |  |
| Likelihood ratio test= 4.74 on 2 df, p=0.09343 |  |  |  |  |  |
| Wald test = 0.75 on 2 df, p=0.6856 |  |  |  |  |  |
| Score (logrank) test = 10.58 on 2 df, p=0.00503 |  |  |  |  |  |

### Comparing dead and alive samples using viral-variant data

We performed variant calling (Module 4) on the liver cancer patients to see if it can add valuable information to the tumor landscape in humans. We then compared the dead and alive samples at the viral-variant level on the 25 HepB patients. For this analysis, the outputs from both Module 1 and 2 were fed into Module 4.

Most of the top variants from filtered human sample (Module 1 + Module 4) and unfiltered human sample (Module 2 + Module 4), were the same. We collated the significant common results (p value <= 0.05) in Table 6 (A) and Table 6 (B). Among these results, we saw several missense and frameshift variants in Gene X of the Hepatitis B genome (nucleotide 1479), Gene P (2573, 2651, 2813), and a region that overlaps Gene P and PreS1 (nucleotides 2990, 2997, 3105, 3156). All these variants were found mutated more in the cases than controls. Other significant common results included variants in Gene C (nucleotide 1979, 2396) and variants in PreS2 region (nucleotide positions 115, 126 and 148) (Table 6 A).

In addition, there were two missense variants that were common among the top results, but not significant (p value = 0.06). They were variants in the X gene of the Hepatitis B genome (nucleotides 1762 and 1764) (Table 6 A).

Among the significant common results to both, were a few variants of the Human endogenous retrovirus K113 complete genome (HERV K113). These include nucleotide positions 7476, 7426 and 8086. These map to frameshift and missense mutations in the putative envelope protein of this virus (Q779_gp1, also called ‘env’) (Table 6 B).

**Table 6:**
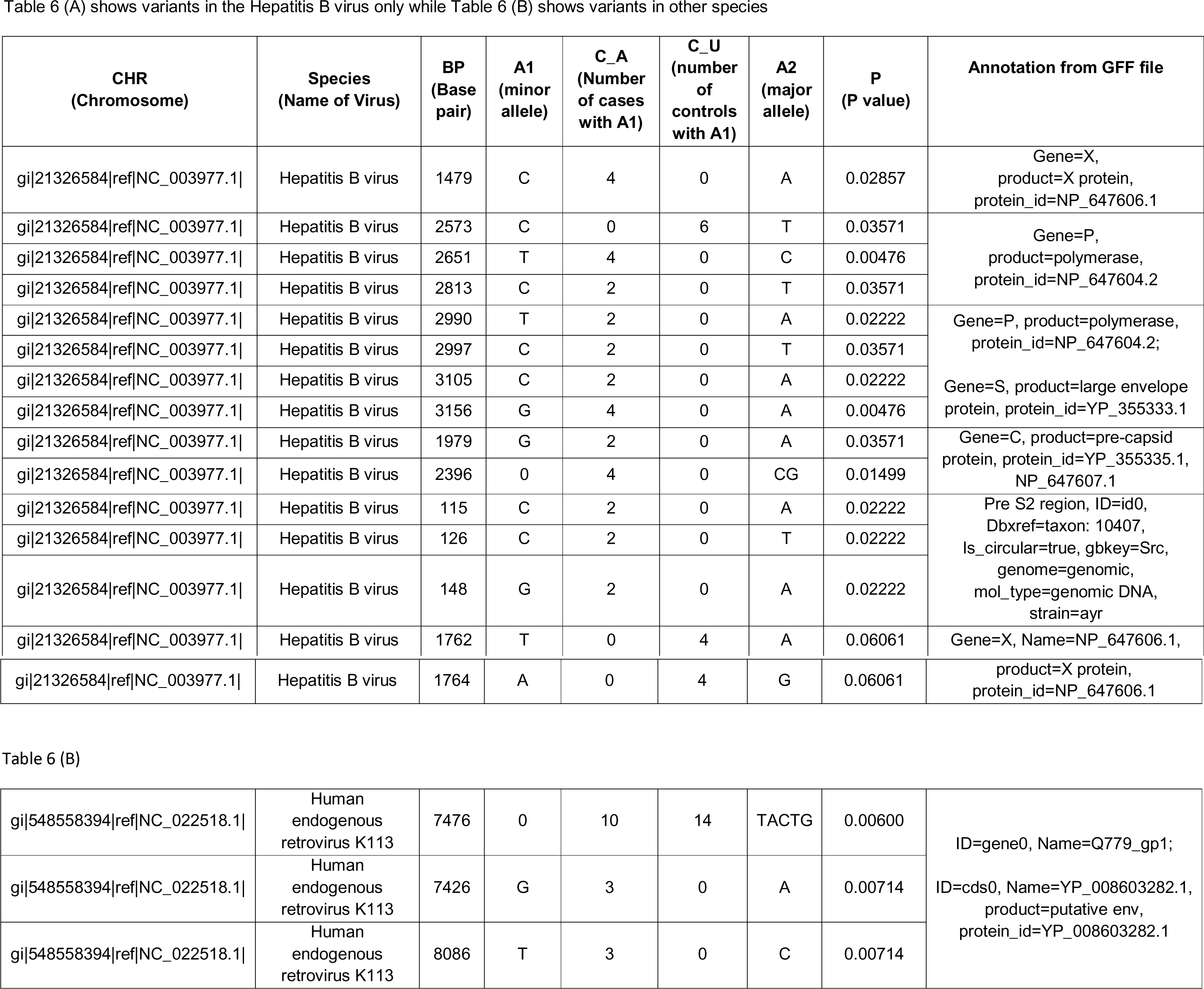
Results of case-control association test applied on the results from viral variant calling. (showing only common results between two possible analysis steps). The table is sorted based on Annotation. Annotation includes gene name, protein name, etc., separated by commas, multiple annotations separated by semi-colon.

## Discussion

### Detection of HPV in cervical cancer patients

The Seven Bridges team used two metagenomic tools Centrifuge [36] and Kraken [37] to detect HPV viruses on the same cohort of TCGA patients [38, 39], and shared the results with us. They used an abundance of 0.02 as a positive viral detection [38, 39]. We compared viGEN with Kraken and Centrifuge in terms of the percentage of samples where the species was detected (Table 7). We can see that the results are in the same range for all three tools.

**Table 7:** Comparing the viral detection ability of viGEN with other tools.

| **Virus Name** | **Tool Name** | **% of samples where the species was found** |
| --- | --- | --- |
| **HPV 16 (Alphapapillomavirus 9)** | Kraken | 58% |
| **HPV 16** | Centrifuge | 55% |
| **HPV 16** | viGEN | 53% |
| **HPV 18** | Kraken | 20% |
| **HPV 18 (Alphapapillomavirus 7)** | Centrifuge | 15% |
| **HPV 18** | viGEN | 13% |
| **HPV 26 (Alphapapillomavirus 5)** | Kraken | 1% |
| **HPV 26** | Centrifuge | 0.3% |
| **HPV 26** | viGEN | 0.3% |

We also estimated the sensitivity and specificity of these tools using the same 22 patients and compared with that of the viGEN pipeline. The Centrifuge tool had a sensitivity of 83% and specificity of 60% for HPV-16 detection; and a sensitivity of 75% and specificity of 94% for HPV-18 detection. The Kraken tool had a sensitivity of 83% and specificity of 20% for HPV-16 detection; and a sensitivity of 75% and specificity of 17% for HPV-18 detection (detailed in Additional File 2). It shows that our viGEN pipeline was able to match the sensitivity and specificity of Centrifuge tool and surpassed that of Kraken (detailed in Additional File 2).

### Additional analysis on liver cancer patients

We used our viGEN pipeline to get genome-level read counts obtained from viruses detected in the RNA of human liver tissue. In our results, HBV was correctly detected in 20% of the samples. This is similar to earlier analyses of TCGA liver cancer cohort study [10, 40, 41], which detected the HBV virus in 23% and 32% (with typically low counts range) of cases respectively.

It has also been reported that the viral gene X (HBx) was the most predominately expressed viral gene in liver cancer samples [40] which is in concordance with our findings where the peak number of reads were observed for gene X region of the HBV genome.

### Comparing dead and alive samples in the liver cancer cohort using viral gene expression data

To get a more detailed overview of the viral landscape, we examined the human RNA-seq data to detect and quantify viral gene expression regions. We then examined the differences between dead and alive samples at the viral-transcript level on the Hepatitis B sub-group (Table 4 (A) and Table 4 (B)).

From the differential expression analyses, the two most informative results were (1) a region of the Hepatitis B genome that produced the HBeAg protein was overexpressed in the dead patients and (2) another region of the Hepatitis B genome that produced HBsAg protein was overexpressed in the alive patients.

Presence of HBeAg or HBcAg is an indicator of active viral replication; this means the person infected with Hepatitis B can likely transmit the virus on to another person. Typically, loss of HBeAg is an indicator of recovery from acute Hepatitis B infection. Active viral replication could allow the virus to persist in infected cells, and increase the risk of disease [42, 43]. So our results, showing that antigens HBeAg and HBcAg were overexpressed in dead patients compared to alive patients makes sense, indicating that these patients never recovered from acute infection.

The results also indicate a higher level of HBsAg in the alive patients compared to the dead patients. The highest levels of HBsAg in the virus are known to occur in the ‘immunotolerant phase’. This pattern is seen in patients who are inactive carriers of the virus i.e. they have the wild type DNA, and the virus has been in the host for so long, that the host does not see the virus as a foreign protein in the body, and hence there’s no immune reaction against the virus. In this phase, there is known to be minimal liver inflammation and low risk of disease progression [44-46]. This could explain why we saw higher level of HBsAg in the alive patients compared to the dead patients.

Also among the significant results were three regions from the Human endogenous retrovirus K113 (HERV K113) genome (with negative log fold change) that were overexpressed in the alive patients. Two of these regions were Sequence-tagged sites (STS) and the third region was in the gag-pro-pol region that has frameshifts. HERV could protect the host from invasion from related viral agents through either retroviral receptor blockade or immune response to the undesirable agent [47].

Overall, we found that our results from viral-gene expression level make biological sense, with much of the results validated through published literature.

### Comparing dead and alive samples in the liver cancer cohort using viral-variant data

We performed variant calling on the viral data to see if it can add valuable information to the tumor landscape in humans. We then compared the dead and alive samples at the viral-variant level on the 25 patients in the Hepatitis B sub-group.

Among the significant results (Table 6A and Table 6B) included variants in Gene C (nucleotide 1979, 2396) and variants in PreS2 region (nucleotide positions 115, 126 and 148). The Gene C region creates the pre-capsid protein, which plays a role in regulating genome replication [48]. The mutation in the 2396 position lies in a known CpG island (ranging from 2215-2490), whose methylation level is significantly correlated with hepatocarcinogenesis [49]. Mutations in PreS2 are associated with persistent HBV infection, and emerge in chronic infections. The PreS1 and PreS2 regions are known to play an essential role in the interaction with immune responses because they contain several epitopes for T or B cells [50].

Mutations in the 1762/1764 positions of the X gene are known to be associated with greater risk of HCC [50] [51], and is independent of serum HBV DNA level [51]. This mutation combination is also known to be associated with hepatitis B related acute-on-chronic liver failure [52]. It is predicted that mutations associated with HCC variants are likely generated during HBV-induced pathogenesis. The A1762T/G1764A combined mutations was shown to be a valuable biomarker in the predicting the risk of HCC [50] [51]; and are often detected about 10 years before the diagnosis of HCC [50].

Among the significant common results to both, were a few variants of the Human endogenous retrovirus K113 complete genome (HERV K113). These variants map to frameshift and missense mutations in the putative envelope protein of this virus (Q779_gp1, also called ‘env’). Studies have shown that this envelope protein mediates infections of cells [53]. HERV K113 is a provirus and is capable of producing intact viral particles [54]. Studies have shown a strong association between HERV-K antibodies and clinical manifestation of disease and therapeutic response [12] [13]. It is hypothesized that retroviral gene products can be ‘reawakened’ when genetic damage occurs through mutations, frameshifts and chromosome breaks. Even though the direct oncogenic effects of HERVs in cancer are yet to be completely understood, it has shown potential as diagnostic or prognostic biomarkers and for immunotherapeutic purposes including vaccines [13].

### Limitations

One limitation of our viGEN pipeline is that it is dependent on sequence information from reference genome. This makes it challenging to detect viral strains where reference sequence information is not known. In the future, we plan to explore de novo assembly when aligning to reference genome.

### Biological significance

In recent years, US regulators approved a viral based cancer therapy [14], proving that the study of viruses in the human transcriptome has biomedical interest, and is paving the way for promising research and new opportunities.

We show that our viGEN pipeline can thus be used on cancer and non-cancer human NGS data to provide additional insights into the biological significance of viral and other types of infection in complex diseases, tumorigeneses and cancer immunology. Detection and characterization of these infectious agents in tumor samples can give us better insights into disease mechanisms and their treatment [2].

## CONCLUSION

With the decreasing costs of NGS analysis, our results show that it is possible to detect viral sequences from whole-transcriptome (RNA-seq) data in humans. Our analysis shows that it is not easy to detect DNA and RNA viruses from tumor tissue, but certainly possible. We were able to not only quantify them at a viral-gene expression level, but also extract variants. Our goal is to facilitate better understanding and gain new insights in the biology of viral presence/infection in actual tumor samples. The results presented in this paper on two case studies are in correspondence with published literature and are a proof of concept of our pipeline.

This pipeline is generalizable, and can be used to examine viruses present in genomic data from other next generation sequencing (NGS) technologies. It can also be used to detect and explore other types of microbes in humans, as long as the sequence information is available from the National Center for Biotechnology Information (NCBI) resources.

This pipeline can thus be used on cancer and non-cancer human NGS data to provide additional insights into the biological significance of viral and other types of infection in complex diseases, tumorigeneses and cancer immunology. We are planning to package this pipeline and make it open source to the bioinformatics community through Bioconductor.

## LIST OF ABBREVIATIONS

HBV: Hepatitis B virus
HCV: Hepatitis C Virus
HERV K113: Human Endogenous Retrovirus K113
TCGA: The Cancer Genome Atlas
HCC: Hepatocellular carcinoma
NAFLD: nonalcoholic fatty liver disease
Hep B: Hepatitis B
Hep C: Hepatitis C
HepB + HepC: coinfected with both Hepatitis B and C virus
HBsAg: Hepatitis B surface antigen
HBeAg: Hepatitis B type e antigen
NGS: next-generation sequencing
RNA-seq: whole transcriptome sequencing
BAM: Binary version of Sequence alignment/map format
CDS: coding sequence
Cox PH: Cox Proportional Hazard
HBx: viral gene X
STS: Sequence-tagged sites
NCBI: National Center for Biotechnology Information
GFF: general-feature-format

## DECLARATIONS

### Availability of data and material

The TCGA liver cancer dataset was used in the analysis and writing of this manuscript. The data can be obtained from https://cancergenome.nih.gov/

Since access to TCGA raw data is controlled access, we could not use this dataset to create a publicly available tutorial. So we looked for publicly available RNA-seq dataset to demonstrate our pipeline with an end-to-end workflow. We chose one sample (SRR1946637) from publicly available liver cancer RNA-seq dataset from NCBI SRA (http://www.ncbi.nlm.nih.gov/bioproject/PRJNA279878). This dataset is also available through EBI SRA (http://www.ebi.ac.uk/ena/data/view/SRR1946637). The dataset consists of 50 Liver cancer patients, and 5 adjacent normal liver tissues. We downloaded the raw reads for one sample, and applied our viGEN pipeline to it. A step-by-step workflow that includes – description of tools, code, intermediate and final analysis results are provided in Github: https://github.com/ICBI/viGEN/.

### Project details

Project name: viGEN

Project home page: https://github.com/ICBI/viGEN/

Operating system(s): The R code is platform independent. The shell scripts can run on Unix, Linux, or iOS environment

Programming language: R, bash/shell

Other requirements: N/A

License: N/A

Any restrictions to use by non-academics: N/A

### Competing interests

The authors do no have any competing interests

### Funding

Not applicable

### Author contributions

KB and YG designed the pipeline. KB and LS implemented the pipeline. KB and YG wrote the manuscript with editorial comments from SM.

## Acknowledgements

Not applicable

## Figures, Tables and Additional Files

**Additional File 1**: viGEN Github tutorial

**Additional File 2:** Detailed results from analysis of TCGA cervical cancer patients

